# Invariant representation of physical stability in the human brain

**DOI:** 10.1101/2021.03.19.385641

**Authors:** R.T. Pramod, M. Cohen, J. Tenenbaum, N. Kanwisher

## Abstract

Successful engagement with the world requires the ability to predict what will happen next. Here we investigate how the brain makes the most basic prediction about the physical world: whether the situation in front of us is stable, and hence likely to stay the same, or unstable, and hence likely to change in the immediate future. Specifically, we ask if judgements of stability can be supported by the kinds of representations that have proven to be highly effective at visual object recognition in both machines and brains, or instead if the ability to determine the physical stability of natural scenes may require generative algorithms that simulate the physics of the world. To find out, we measured responses in both convolutional neural networks (CNNs) and the brain (using fMRI) to natural images of physically stable versus unstable scenarios. We find no evidence for generalizable representations of physical stability in either standard CNNs trained on visual object and scene classification (ImageNet), or in the human ventral visual pathway, which has long been implicated in the same process. However, in fronto-parietal regions previously implicated in intuitive physical reasoning we find both scenario-invariant representations of physical stability, and higher univariate responses to unstable than stable scenes. These results demonstrate abstract representations of physical stability in the dorsal but not ventral pathway, consistent with the hypothesis that the computations underlying stability entail not just pattern classification but forward physical simulation.

## Introduction

Visual percepts are imbued with possibility. We see not just a dog pointing motionless at a rabbit, but a dog about to give chase; not just a wineglass near the edge of a table but a wineglass about to smash on the floor; not just a cup filled to the brim with coffee, but a cup at risk of staining the white tablecloth beneath. This ability to see not only the current situation in front of us, but what might happen next, is essential for planning any action. Although some of our predictions concern other people and what they will think and do, many of our predictions concern the physical world around us. The most basic prediction we make about the physical world is whether it is stable, and hence unlikely to change in the near future, or unstable, and likely to change. Here we ask whether the brain computes physical stability via simple pattern classification, such as that thought to be conducted in the ventral visual pathway, or whether instead we determine stability by running a kind of simulation in our head to determine what, if anything, will happen next.

What kind of representations might be required for determining the physical stability of a scene? Computational work has shown that convolutional neural networks (CNNs) trained on ImageNet learn powerful representations that support not only object recognition, but multiple other tasks^1–4^. Indeed, several studies have claimed that these networks can accurately predict scene stability^5–7^. But CNNs have so far been tested only within very narrow domains (e.g., block towers with several cubes of different colors, in Fig 1a, left column) and it is not clear they would successfully predict future states of complex natural images (e.g., Fig 1a. middle two columns). Indeed, CNN models of stability trained on towers of blocks have not been shown to generalize even to other kinds of block configurations, with different numbers, sizes, shapes, colors and configurations of blocks than those encountered during training. A *general* ability to predict what will happen next in physical scenarios may instead require a richer representational of the physical properties of the scene that support forward simulations like those in the physics engines of video games^8,9^. Behavioral evidence from humans is conflicting here. On the one hand, human observers can perceive the stability of a tower of blocks extremely rapidly and pre-attentively^10^, and these findings have been taken to argue for a simple feed-forward rather than a simulation process. On the other hand, human judgements of the stability of a tower of blocks are well modeled by approximate probabilistic simulation in game-style physics engines^8,9,11,12^, which better fit the human pattern of errors than CNNs do^11^. Perhaps most importantly, humans can easily judge the physical stability of a wide variety of novel real-world images, which is likely to require a more abstract and generalizable representation of physical properties of the scene. Here we test whether representations that support fast feedforward visual object and scene recognition in machines (i.e., CNNs trained on ImageNet) and brains (i.e., the ventral visual pathway) can support discrimination of the physical stability of complex natural scenes, or whether different kinds of computations – in particular, those that explicitly model dynamic physical properties of the scene, perhaps through a kind of simulation mechanism --may be required to determine physical stability in both machines and brains.

**Figure 1.**
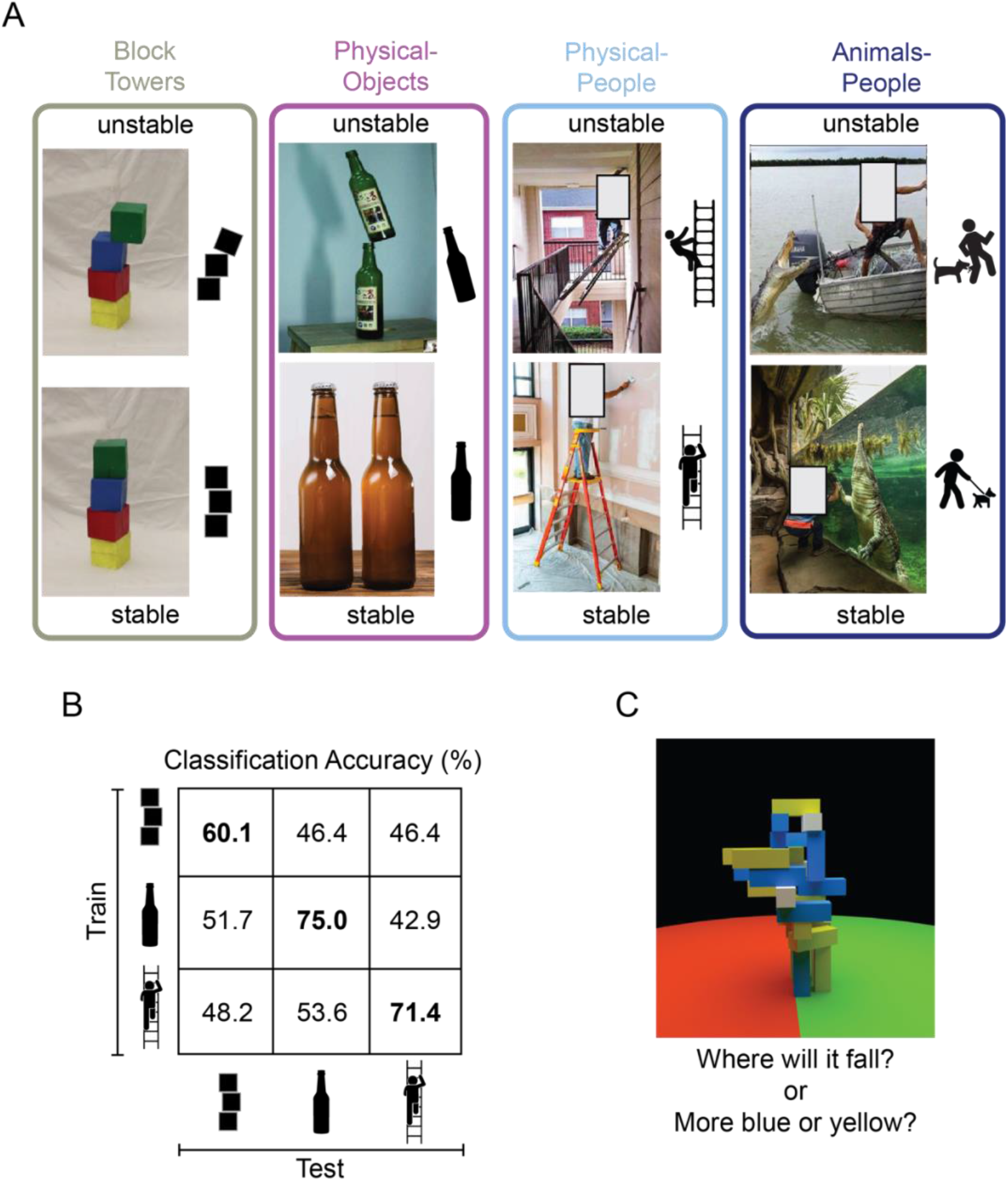
Design of Studies Conducted on Machines and Brains. (A) Image Sets depicting (i) unstable vs stable block towers (*left*), (ii) objects in unstable versus stable configurations (*middle left; the Physical-Objects set*), (iii) people in physically unstable versus stable situations (*middle right, the Physical-People set*), and (iv) people in perilous (“unstable”) versus relatively safe (“stable”) situations with respect to animals (*right*, the *Animals-People* set). Note that by “physically unstable” we refer to situations that are currently unchanging, but that would likely change dramatically after a small perturbation like a nudge or a puff of wind. The former three scenarios were used to study the generalizability of feature representations in a deep convolutional neural network (CNN) and the latter three scenarios were used in the human fMRI experiment. (B) Accuracy of a linear support vector machine (SVM) classifier trained to distinguish between CNN feature patterns for unstable versus stable conditions, as a function of Test Set. Accuracy within a scenario (i.e., train and test on the same scenario) was computed using a 4-fold cross-validation procedure. (C) A functional localizer task contrasted physical versus non-physical judgements on visually identical movie stimuli (adapted from Fischer et al, 2016). During the localizer task, subjects viewed movies of block towers from a camera viewpoint that panned 360 degrees around the tower, and either answered “where will it fall?” (physical) or “does the tower have more blue or yellow blocks?” (non-physical).

To investigate this question, we first curated a set of natural images depicting real-world scenarios of objects in physically stable versus unstable configurations (the “Physical-Objects” set, Figure 1A), as well as a generalization set of images of people in physically stable versus unstable situations (the “Physical-People” set, Figure 1A). We then presented these stimuli to CNNs pretrained on ImageNet classification and asked if the resulting representations are sufficient to discriminate the physical stability of the scene in a fashion that generalizes across image sets. Next, we presented the same stimuli to human participants being scanned with fMRI and asked the same question of the resulting neural responses in two key brain regions. One such region is the ventral visual pathway, which is widely thought to conduct the core computations underlying visual object recognition, and which is well modelled by CNNs trained on ImageNet^13,14^. Thus, if representations optimized for object recognition suffice for determining the physical stability of novel scenes, we would expect to find this ability in the ventral visual pathway. However, an alternative hypothesis is that this information will be found instead in a set of parietal and frontal regions that have been shown to be engaged when people predict what will happen next in physical (more than social) scenarios^15^, and hold information about physical variables such as mass, invariant to the scenario that revealed that mass^16^. We have hypothesized^15^ that these regions (referred to here as the “Physics Network”) may contain a generative model of dynamic physical scenes capable of forward simulation, which can be used for prediction, planning, and many other tasks. Thus, if determining the stability of a scene requires an abstract representation of the physical properties of the scene and the ability run forward simulations on those representations, then the Physics Network would be a more likely locus of these representations than the ventral visual pathway.

Finally, we further conjectured that if perception of physical stability is determined by the operation of a forward simulation mechanism akin to a generative physics engine, then we might find higher neural activity in the Physics Network for physically unstable than stable situations because there is more to simulate in an unstable situation. That is, simulations of unstable scenes where objects are expected to move require the construction and updating of dynamic representations that unfold over time, whereas simulations of stable scenes, where objects are expected to remain stationary, do not require such dynamically updated representations. Importantly, participants were never asked to judge or think about the physical stability of the scenario they were viewing, which enabled us to ask whether this information is extracted even when not required by the task^17^.

## Results

### Feed-forward convolutional neural networks trained on ImageNet do not have a generalizable representation of physical stability

Previous studies have shown that visual features extracted by feed-forward deep convolutional neural networks (like AlexNet^18^, VGG^7^, and ResNet^5,6^) can distinguish between stable and unstable towers. Although these features produced above-chance classification performance within a scenario (i.e., train on towers with 2 blocks and test on towers with 4 blocks), they have not been tested on very different scenarios and hence their ability to represent an abstract and generalizable notion of physical stability remains unknown. Significant generalizability of these features to novel image sets would provide some evidence that physical stability can be inferred from a simpler mechanism of visual feature extraction in pattern classification systems rather than running forward simulations of the kind used in game physics engines.

To test this question, we extracted features for our stable and unstable images from the final fully connected layer (*fc1000*, the layer immediately preceding the softmax probability layer) of ResNet-50 – a feedforward deep convolutional neural network (CNN) trained on ImageNet dataset for object recognition^19^. A shallower variant of this network has been previously used for stability classification^5,6^. Within each scenario (Block Towers, Physical-Objects, and Physical-People, Figure 1A), we trained a linear Support Vector Machine (SVM) classifier to distinguish between feature patterns for stable and unstable images and tested it on held-out images using 4-fold cross-validation. These analyses replicated previous results by showing above chance classification accuracy within the Block Towers scenario and further found higher classification performance within our two new natural image sets (60.1%, 75% and 71.4% for Block Towers, Physical-Objects, and Physical-People scenarios respectively; diagonal cells in Figure 1B). However, this performance did not generalize across scenarios: accuracy was near chance for all cross-validation tests in which the SVM classifier was trained on one scenario and tested on a different scenario (i.e., off-diagonal cells in Figure 1B). Thus, visual features from this feedforward deep convolutional neural network do not carry abstract and generalizable information about physical stability. We also found similar results with a VGG-16 network trained on ImageNet.

These results indicate that the ImageNet-trained CNNs previously claimed to support judgements of stability cannot do so in a fashion that generalizes to novel image sets. This result further suggests that brain regions thought to be optimized (through development and/or evolution) for similar tasks (Ventral Temporal Cortex) might similarly fail to contain scenario-invariant information about physical stability, and that the human ability to perform this task may result from different kinds of representations elsewhere in the brain, such as the fronto-parietal cortical regions previously implicated in physical inference. We test these hypotheses next.

### A generalizable representation of physical stability in cortical regions previously implicated in intuitive physical inference, but not in the ventral temporal cortex

We first identified functional regions of interest (fROI) in each participant individually, including Ventral Temporal Cortex (VTC) and the candidate Physics Network in the frontal and parietal lobes, by intersecting anatomical constraint parcels for these regions with the relevant functional activations (see Figure 2A and Methods). All participants had significant voxels within both VTC and physics parcels (number of voxels per participant, mean ± sem = 350.5 ± 68.7 for VTC ROIs and mean ± sem = 311.3 ± 69.8 for physics ROIs).

**Figure 2.**
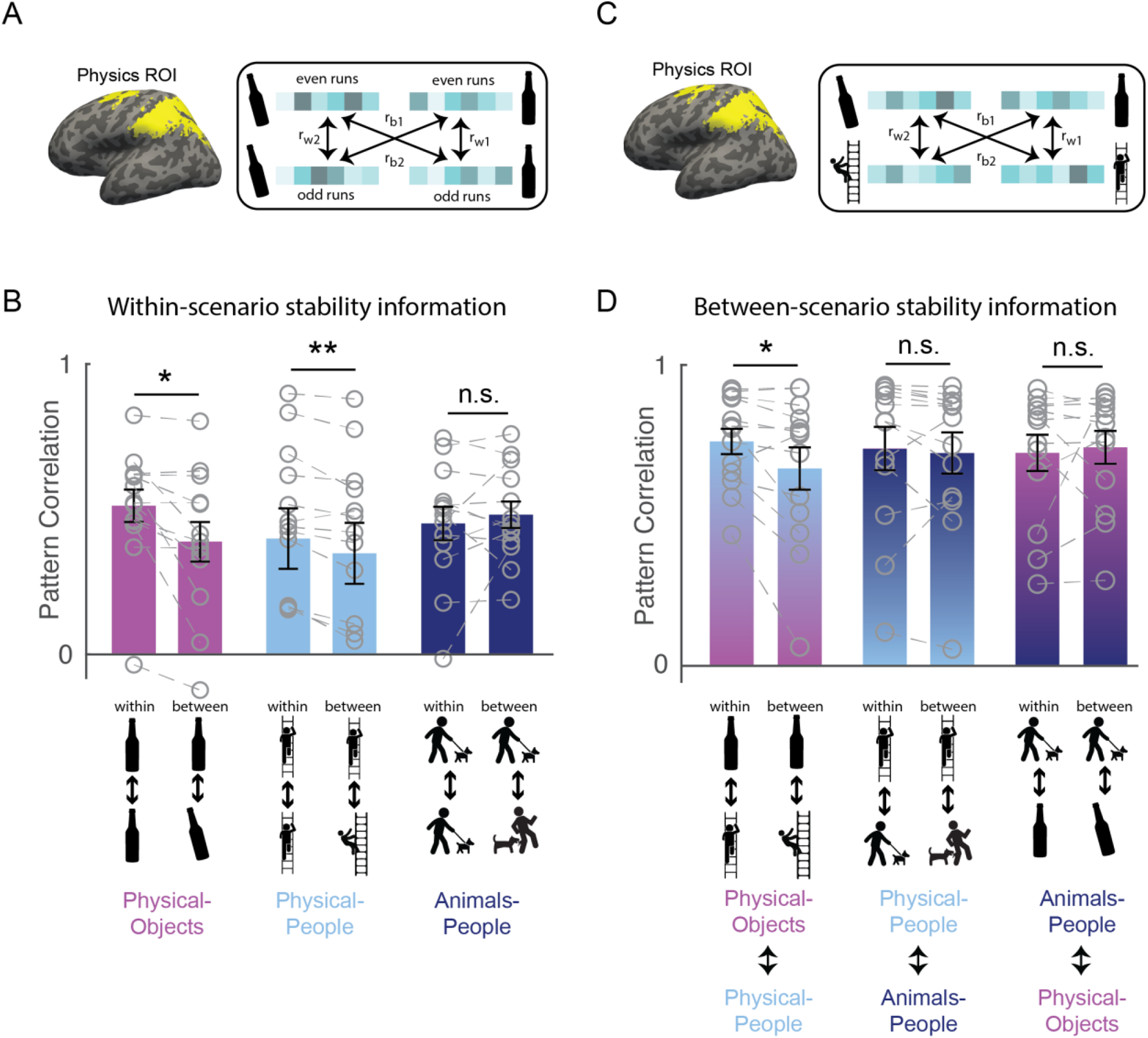
(A) Functionally defined regions of interest implicated in intuitive physical inference (the Physics Network) were defined in each participant individually by intersecting the activation from a localizer task with anatomical constraint parcels in the frontal and parietal lobes shown in yellow (see Methods). Patterns of activity across voxels in these fROIs were extracted separately in each participant for each combination of even and odd runs, unstable and stable conditions, and scenario (Physical-Objects, Physical-People, and Animals-People). Following standard practice^20^, correlations between even and odd runs in the pattern of response across voxels were computed both within stability conditions (rw1 = stable even to stable odd, and rw2 = unstable even to unstable odd), and between stability conditions (rb1 = stable even to unstable odd, and rb2 = unstable even to stable odd), for each of the three scenario types. (B) Bar plot showing average pattern correlations *within* and *between* conditions with paired t-tests done after Fisher transformation, for each scenario type. (C) is similar to (A) but the within and between stability pattern correlations were computed across scenario types (rw1 and rw2 indicate pattern correlations computed within stable or *within* unstable conditions across scenario, and rb1 and rb2 indicate pattern correlations computed *between* stable and unstable conditions across scenario). (D) is same as in (B) but for pattern correlations computed across scenarios. Note that the across-scenario correlations in (D) are overall higher than within scenario correlations in (C) because the data was not split into odd and even runs for the across-scenario case. Grey circles and the corresponding connecting lines denote individual subject”s data. Error bars indicate standard error of mean across subjects. ** and * indicate statistically significant effect at p <= 0.005 and p <= 0.05 respectively, and n.s. indicates no statistically significant effect.

#### Within-Scenario Stability Information

To test whether VTC and Physics Network contain information about physical stability, we measured the fMRI response of each voxel of the fROI in each participant, separately for odd and even runs in each of the six stimulus conditions of the main experiment. We then used multi-voxel pattern analysis^20^ to test whether physical stability could be decoded in each fROI. Specifically, across all voxels in each fROI we computed the correlation between the response patterns *within* stability conditions (stable even to stable odd, and unstable even to unstable odd) and *between* stability conditions (stable even to unstable odd, and stable odd to unstable even) for each of the scenarios separately (see Figure 2A and Methods). For VTC, we found that the *within* condition pattern correlations were significantly higher than the *between* condition pattern correlations for both Physical-Object and Physical-People scenarios (see Table 1), indicating the presence of information about stability for each scenario type. However, pattern information did not distinguish perilous (or “unstable”) from non-perilous (“stable”) situations in the Animals-People scenarios (see Table 1 for stats). Further, this difference in stability information across scenario types was itself significant, as revealed by a significant interaction between scenario type and within versus between correlations in an ANOVA (F_1,12_ = 72.2, p = 0.000002). The Physics Network showed the same pattern of results (Figure 2B and Table 1), with significantly higher correlations within than between stability for both the Physical-Object and Physical-People scenarios but not the Animals-People scenarios, and a significant ANOVA interaction between scenario type and within versus between correlations (F_1,12_ = 22.9, p = 0.0004). Thus, both VTC and the Physics Network contain information about physical (but not animate) stability when analyzed within scenarios.

**Table 1:**
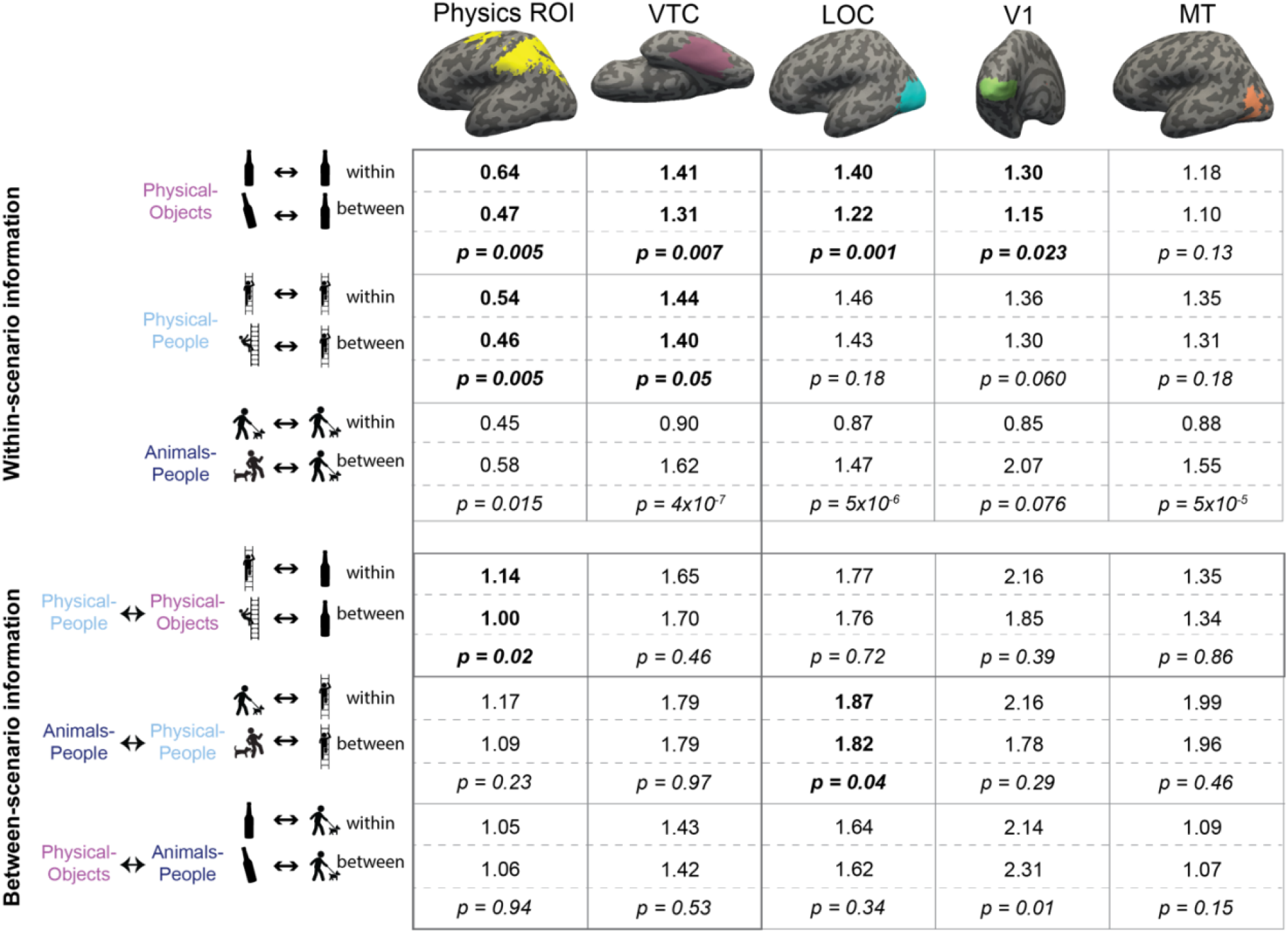
Multi-voxel pattern correlation analysis in all ROIs. Each cell shows the average Fisher transformed correlation within and between conditions, along with the p-value for a paired t-test comparing the two sets of values. Each column includes the results from one fROI. The top three rows contain results for the analyses within scenario type and the bottom three rows show results for the pattern correlation analysis across scenarios. Significantly higher correlations for within than between conditions are highlighted in bold. The row depicting the crucial cross-decoding result with significant generalization only in the Physics Network ROI is highlighted with a darker bounding box.

#### Between-Scenario Stability Information

However, our strong prediction was that any brain region that represents abstract physical information should show patterns of response that generalize across physical scenarios (Physical-Objects and Physical-People). To test this, we again used multi-voxel pattern correlation analysis to compute *within*-condition (unstable-unstable and stable-stable) and *between*-condition (unstable-stable and vice versa) correlations, but this time across scenario types (Figure 2C). Indeed, we found that in the Physics Network (and not in VTC), the within condition pattern correlations are significantly higher than between condition pattern correlations only across Physical-Objects and Physical-People scenarios, but not across Physical and Animate scenarios (Figure 2D and Table 1). Further, we found a significant interaction between fROI (physics versus VTC) and between-scenario stability information (within versus between correlations), indicating stronger generalization across scenarios in the Physics Network compared to the VTC fROI (F_1,12_ = 6.29, p = 0.027). Thus, the Physics Network (and *not* VTC) holds information about physical stability that generalizes from scenarios with only inanimate objects to scenarios in which people are in physically unstable situations, but not to scenarios in which people are in peril from animals.

#### Eye Movement and Attention Controls

Could these findings in the Physics Network simply be due to differential eye movements across different experimental conditions (despite the instructions to fixate), or differential attention? To explore these possibilities, we quantified eye movements collected in the scanner during the main experiment in a subset of participants (n = 6) for each of the six stimulus conditions by extracting the average x and y co-ordinates of eye position, as well as the number, duration and amplitude of eye movements. None of these quantities showed a significant difference between stable and unstable conditions in any of the scenarios except for saccade amplitude in the “Physical-People” scenario (p = 0.028, Supplementary Table 1). Moreover, we found no significant interaction of stability with scenario type for any of the eye movement quantities using separate ANOVAs (p > 0.1). Second, analysis of subjective ratings by a separate set of subjects (see Methods) of “*interestingness*” of our stimuli (which we used as a proxy for how attention-grabbing the stimuli were) revealed significantly higher ratings for unstable over stable conditions in all three scenarios (p < 0.001 for a paired t-test on average ratings across subjects; Supplementary Table 1, last column). However, we found no interaction between stability and scenario type in the ANOVA (p = 0.23), implying that the difference in ratings between stability conditions did not vary between physical and animate scenarios. Thus, the distinctive and generalizable representation of physical stability we found in the Physics Network are unlikely to be due to differential eye movements or attention.

#### Analysis of Subregions of the Physics fROIs

The Physics Network considered here include regions in both the parietal and frontal lobes in both hemispheres. Evidence that the scenario-invariant representation of physical stability reported above is not restricted to a subset of these regions comes from separate analyses of the left parietal, left frontal, right parietal, and right frontal fROIs in each participant. An ANOVA analyzing the generalization of stability information (between versus within condition pattern correlations across Physical-Objects and Physical-People; F_1,12_ = 6.11, p = 0.03 for the main effect) did not find a significant interaction of stability information with either hemisphere (left vs. right; F_1,12_ = 2.75, p = 0.12) or lobe (parietal vs. frontal; F_1,12_ = 0.24, p = 0.63; Supplementary Table 2).

### Other visual regions do not have a generalizable representation of physical stability

In the previous section, we showed that the Physics Network in the fronto-parietal cortices, but not VTC, has a representation of physical stability that generalizes across scenario. Is this abstract representation also found in other visual regions? We functionally defined two other visual regions: V1 and LOC (see Methods) and performed multi-voxel pattern correlation analyses as before. In both regions, we found significant decoding of stability only in Physical-Objects scenario but no cross-decoding of stability across physical scenarios (Table 1). In addition, the interaction of this cross-decoding effect with fROI was significant in separate ANOVAs contrasting the Physics Network with V1 (F_1,12_ = 6.31, p = 0.027), and LOC (F_1,12_ = 5.63, p = 0.035). Thus, a generalizable representation of physical stability is apparently a distinctive property of the Physics Network in the parietal and frontal lobes and is not a widespread property of visual cortex.

### Higher Mean Responses to Physically Unstable than Stable Scenes in Physics fROI

Previous studies have proposed that human intuitive physical reasoning, including inferences about physical stability, can be explained by a model that performs probabilistic simulations of the future states of the physical world^8,11^. Do the candidate physics regions in our brain perform this forward simulation of what will happen next? Some evidence for this idea comes from the higher response of these regions during physical prediction than color-judgement tasks used in our localizer and in the original fMRI study that used this task^15^. However, neural activity in that contrast could simply reflect the process of building a mental model of the physical scene (including object properties and relationships), not predicting or simulating what would happen next. Here we reasoned that if the candidate physics regions are engaged automatically in simulating what will happen next, they should show a higher mean response when viewing physically unstable scenes (because there is more to simulate) than stable scenes (where nothing is predicted to happen).

We tested this prediction by comparing the average fMRI response for unstable and stable conditions in each of the three scenario types, in the Physics Network (defined in the same way as in the previous analysis). As predicted, the physically unstable condition showed a significantly greater response compared to the physically stable condition in both “Physical-Objects” (p = 0.02 for a paired t-test on average response across subjects) and “Physical-People” scenarios (p = 0.04 for a paired t-test on average response across subjects; Figure 3 & Table 2 left column), but not the “Animals-People” scenario (p = 0.66 for a paired t-test on average response across subjects; Figure 3 & Table 2). As before, we also checked whether this trend is largely driven by voxels in one of the hemispheres (right/left) or one of the lobes (parietal/frontal) by performing an ANOVA. We found a significant interaction between stability (unstable versus stable) and lobe (F_1,12_ = 8.28, p = 0.014) but not hemisphere (F_1,12_ = 3.04, p = 0.11) for the “Physical-Objects” scenarios, and the same pattern for the “Physical-People” scenario (stability x lobe: F_1,12_ = 8.53, p = 0.013; stability x hemisphere: F_1,12_ = 0.34, p = 0.57). Post-hoc analysis revealed that this interaction effect was largely driven by the univariate difference in the parietal but not frontal lobe (Supplementary Table 3).

**Table 2:**
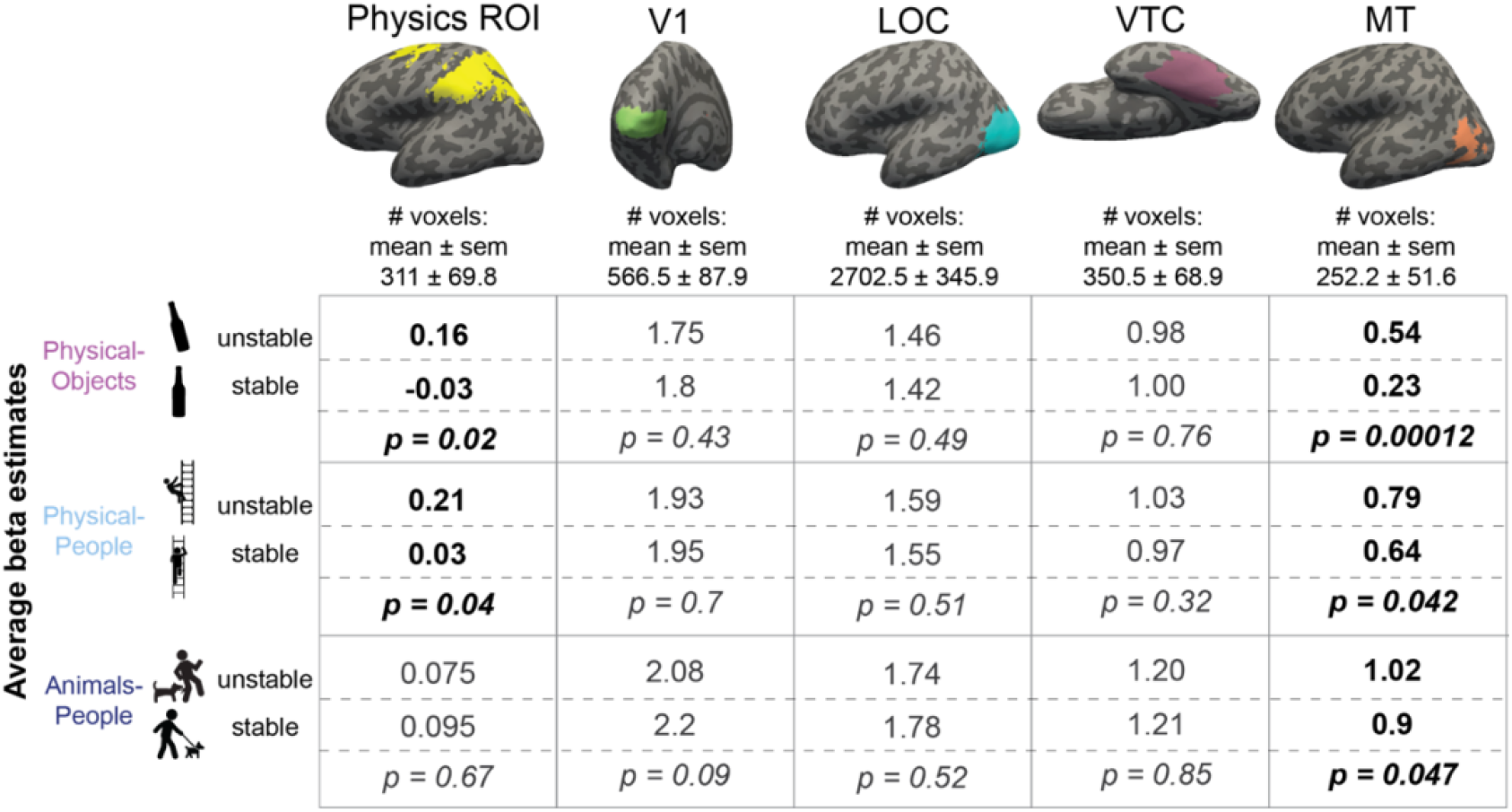
Average beta values for unstable and stable conditions in each of the scenarios. Columns represent different fROIs. Each cell shows average GLM estimated beta values for unstable and stable conditions along with the p-value for a paired t-test comparing the two sets of values. Scenarios showing significantly higher response to unstable scenes compared to stable scenes are highlighted in bold in each column.

**Figure 3.**
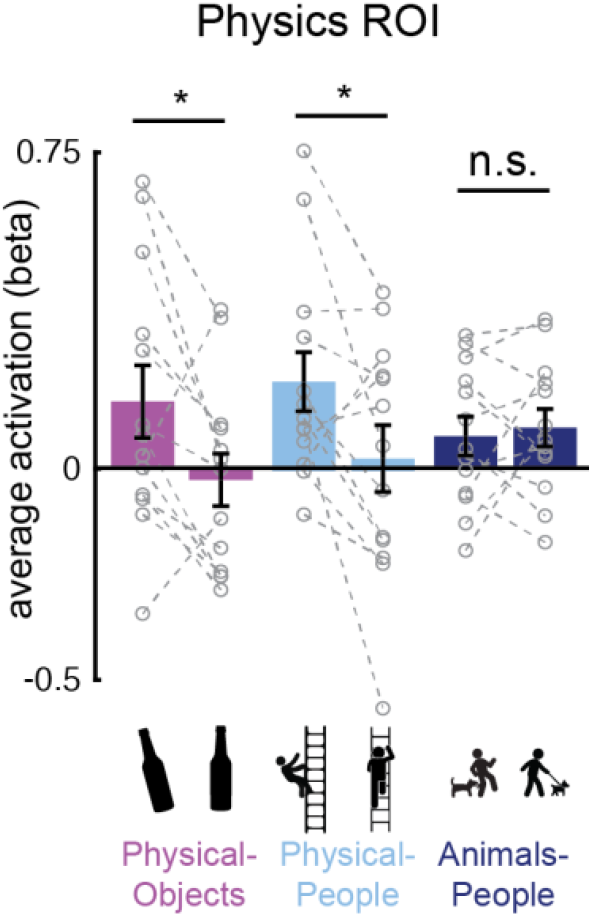
Bar plot showing the average activation (GLM beta estimates) in the physics fROI for both stable and unstable conditions in each of the three scenario types (Physical-Objects, Physical-People, Animals-People). Grey circles and the corresponding connecting lines indicate individual subject”s data. Error bars indicate standard error of mean across subjects. * indicates significant effect at p < 0.05 and n.s. indicates no statistically significant effect.

According to our hypothesis, it is the automatic simulation occurring in the Physics Network that leads to the higher response for unstable than stable conditions. As such we predicted that we would not see this same effect in VTC and other visual regions (V1 and LOC) that are not engaged in physical simulation. Indeed, as shown in Table 2, we did not. Furthermore, we found a significant interaction of stability with region (physics fROI vs. visual fROI) for the “Physical-Objects” (F_1,12_ = 13.04, p = 0.0035 for V1; F_1,12_ = 8.27, p = 0.014 for VTC) and the “Physical-People” scenarios (F_1,12_ = 11.14, p = 0.0059 for V1; F_1,12_ = 7.83, p = 0.016 for LOC).

Thus, physically unstable scenes evoke stronger responses than stable scenes in the fronto-parietal Physics Network, but not elsewhere, consistent with the hypothesis that these regions are engaged in running forward simulations of what will happen next.

#### Higher responses to Any Instability in Visual Motion Area MT

Finally, we predicted that visual motion area MT might show higher responses for all three forms of instability, whether physical (with objects or people), or animate. Our rationale was that because MT is implicated in motion processing in general, any forward simulation involving motion, whether physical or animate, could activate this region. Indeed, that is exactly what we saw. As shown in Tables 1 & 2 (last column), we found significantly higher responses in MT for unstable/perilous than stable/safe scenes for all three types of scenarios without any significant stability information in the pattern activations. This increased response to conditions with predicted motion is reminiscent of previous reports showing greater response in MT for static images with implied motion^21^. The critical difference, however, is that whereas the earlier study reported higher responses in MT for static images depicting motion events happening at the moment the photograph was taken (implied motion) the current study shows activation of MT for motion that is only *predicted* (in the Physical-Objects and Physical-People scenarios).

## Discussion

Here we report that fronto-parietal cortical regions previously implicated in intuitive physical inference contain abstract information about physical stability, but representations in the ventral visual object recognition pathway, and those in feedforward convolutional neural networks trained on object recognition, do not. These results indicate that representations in systems that are highly effective at invariant object recognition do not automatically support the general ability to distinguish physically stable from unstable scenes. Instead, this ability in humans is supported by a different system in the dorsal visual pathway that has been previously implicated in intuitive physical inference. This Physics Network (but not the ventral pathway) further shows a higher univariate response to unstable than stable scenes, as predicted if it performs forward simulations of what will happen next. Control analyses confirmed that neither pattern nor univariate information about physical stability in the Physics Network can be accounted for by low-level visual features, differential eye movements or attention. Taken together, these results suggest that the human brain represents the physical world not via simple pattern classification but instead by building a model of the physical world that supports prediction via forward simulation.

Our study builds upon earlier work that has implicated regions in the parietal and frontal lobes in intuitive physical inference. Across multiple studies, similar fronto-parietal regions have been shown to a) respond more during physics-based tasks compared to color-based or social prediction tasks^15^, b) represent physical concepts in verbal stimul^22^, and c) contain scenario-invariant representations of object mass^16^. Intriguingly, the regions activated during physical inference overlap with regions shown previously to be engaged during visually-guided action^23^ and tool use^24–26^, perhaps because these tasks also require a representation of the physical world. Indeed physical simulation has been proposed as a crucial component in models for human-like robotic action planning^27–29^ and flexible tool use^30^; plausibly the same brain mechanisms could have arisen or adapted to support all of these computations. Note however that these same regions also overlap with the “multiple demand” system^15^, and are unlikely to be selectively engaged in *only* physical inference. In the present study, we strengthen evidence that these regions are engaged in intuitive physical inference by showing that they carry a new kind of physical information: the dynamic physical stability of objects and people in a scene. Interestingly, this information is present when participants simply view images of physical scenes, even though they are not asked to judge stability.

More importantly, our work speaks not only to which brain regions are implicated in intuitive physical inference, but what kinds of computations these inferences entail in both minds and machines. In particular, it is a matter of active current debate in AI whether features extracted by feedforward computations in networks trained on visual object classification will suffice for physical inference, or whether richer internal models that support forward simulation are required. Some work has shown that CNNs can learn from extensive labeled datasets to infer the stability of block towers^5,6^ and predict future outcomes^5–7^. But these CNNs have so far been tested only on specific tasks and stimuli, and we show here that they do not generalize across scenarios for the task of determining physical stability. Instead, it has been proposed that a general ability to predict what will happen next in physical scenarios will require a more structured representation of the physical world that will support forward simulation.^8,9^ A parallel debate is raging in cognitive science^5,6,8,10,17,31–34,35^, between those who argue that because human physical inferences occur rapidly^10^ and pre-attentively^10^ they are computed by something like a pattern recognition process, versus those who argue that human and primate physical inference behavior is best accounted for by mental simulation^8,9,11,12,36^. Three lines of evidence from the present study support the simulation view. First, we find generalizable representations of physical stability in the brain that we do not find in CNNs. Second, these abstract representations of stability are not found in the ventral visual pathway, which is thought to conduct pattern classification and is well modeled by CNNs, but rather in the dorsal pathway, previously implicated in intuitive physical inference. Third, we find a higher univariate response in this Physics Network for unstable scenes, where there is more to simulate, than the stable scenes, where nothing is predicted to happen next.

We therefore hypothesize that visual information represented in the ventral visual cortex is used by the Physics Network for efficient inference of physical properties, and this representation is in turn used for forward simulation of what will happen next. This idea has been recently proposed as an integrated computational model that uses visual representations from a deep-learning-based inverse graphics model to initialize simulations in a physics-engine-based generative model of object dynamics, which then can be used to perceive, predict, reason about and plan with physical objects^37–39^. This class of models with flexible, object-centric representations and the ability to learn from realistic visual inputs^40,41^ should be able to make predictions of physical stability on the realistic stimuli used in our experiment and also form the basis for neurally-mappable encoding models of the candidate physics regions. The question of how such a model is instantiated, if at all, in the brain remains unanswered and provides a fertile avenue for future exploration.

Many questions remain. First, physical stability is just one of many aspects of physical scene understanding. Future investigations can explore whether the Physics Network also represents other physical properties of objects (like friction and elasticity), relational attributes (like support, containment, attachment), and physical forces and events. Second, if indeed the Physics Network is conducting forward simulations, when exactly does it run and how detailed are its simulations? According to one hypothesis, our mental physics engine compresses the rich details in our visual world into a relatively small number of individual objects and associated events in order to efficiently generate a reasonable approximation of the scene at the spatial and temporal scales relevant to human perception and action^9^. These abstractions may also enable us to run simulations faster than real-time with compressed timescales (like hippocampal replay^42^), enabling us to make rapid and accurate predictions of the consequences of multiple actions under consideration, including our ability to make fast and automatic physical inference^10,17^. Third, is the same neural machinery underlying simulation of the external physical world also recruited when we consider the consequences of our own actions? Answering this question would help elucidate how action planning and tool use are related to the neural system for physical inference, given that much of the Physics Network lies adjacent to or overlaps with brain regions engaged in action planning and tool use^24–26,43,44^.

## Acknowledgements

This work was supported by NIH grant DP1HD091947 to N.K, a US/UK ONR MURI project (Understanding Scenes and Events through Joint Parsing, Cognitive Reasoning and Lifelong Learning), and National Science Foundation Science and Technology Center for Brains, Minds, and Machines Grant CCF-1231216. The Athinoula A. Martinos Imaging Center at MIT is supported by the NIH Shared instrumentation grant #S10OD021569. We thank Kirsten Lydic for helping with data collection, David Beeler for helping with fMRI data pre-processing, and Ilker Yildirim, Kevin Smith, and members of the Kanwisher Lab for helpful discussions.

## Materials and methods

### Participants

13 subjects (ages 21-34; 6 female) participated in the experiment. All participants were right-handed and had normal or corrected-to-normal vision. Before participating in the experiment, all subjects gave informed consent to the experimental protocol approved by the Massachusetts Institute of Technology (MIT) Committee on the Use of Humans as Experimental Subjects. The study was conducted in compliance with all the relevant ethical guidelines and regulations for work with human participants.

### Stimuli

All images were chosen to belong to six different experimental conditions divided into three scenarios with two conditions each (see Figure 1B for examples): objects in stable or unstable conditions (“Physical-Objects”); people in physically stable or unstable conditions (“Physical-People”); and, people with animals in perilous (unstable) or relatively safe (stable) conditions (“Animals-People”). All images were scaled so that the longer dimension measured 8° on the projector screen places ∼136cm from the participant.

#### Screening using deep neural networks

We chose images from each of these six experimental conditions such that stability decoding accuracy was close to chance both within the scenario and also across scenarios on features extracted from the initial layers of a deep convolutional neural network. The main goal of this selection process was to screen images to minimize potential confounds of low-level visual features on stability decoding. First, we rescaled and padded each image with pixels of zero brightness to obtain images measuring 224 x 224 pixels. Then, we extracted features from the first pooling layer (“pool_1”) of a feedforward convolutional network, VGG-16, trained on ImageNet object classification task. We then trained separate linear SVM classifiers with 4-fold cross-validation for each scenario separately to distinguish between stable and unstable images. The classification accuracies were close to chance (= 50%) for all three scenarios (% accuracy: 42.9%, 53.6% and 42.9% for Physical-Objects, Physical-People and Animals-People scenarios respectively). We then tested each classifier on the remaining two scenarios to quantify cross-decoding of stability across scenarios. Here also, we found close to chance (= 50%) cross-decoding performance (average % accuracy = 51.8% for Physical-Objects vs. Physical-People, 44.6% for Physical-Objects vs. Animals-People, and, 41.1% for Physical-People vs. Animals-People).

#### Interestingness ratings

In order to minimize the influence of differential attention or interest on our results, we set out to quantify how interesting or attention-grabbing our stimuli are. We ran a behavioral experiment on 11 subjects (9 of them had previously participated in the fMRI part of this study) where we asked them to rate how *interesting* they found an image to be on a scale of 1-5 (1 – least interesting and 5 – most interesting). As expected, subjects found unstable condition to be more interesting than stable condition across scenarios (Supplementary Table 1, last column). However, we found that this difference did not significantly interact with scenario using an ANOVA (p = 0.23).

### Experimental design

#### Physics ROI localizer

Each participant performed 2 runs of an “intuitive physics” fMRI localizer task previously used to functionally define the fronto-parietal physics engine in the brain^15,16^. In this task, subjects viewed short movies (∼ 6s) depicting unstable towers made of blue, yellow and white blocks (see Figure 1A) created using Blender 2.70 (Blender Foundation). The tower was centered on a floor that was colored green on one half and red on the other half such that it would topple towards one of the halves if gravity were to take effect. Throughout the movie, the tower remained stationary while the camera panned 360° to reveal different views of the tower. Subjects viewed these movies and were instructed to report whether more blocks would come to rest on the red or green half of the floor (“physics” task), or whether there are more blue or yellow blocks in the tower (“color” task).

Each run of this localizer task consisted of 23 18s blocks: 3 fixation-only blocks, 10 blocks each of the physics and the color task. Each 6s movie was preceded by a text instruction displayed on the screen for 1s which read either “where will it fall?” (“physics” task) or “more blue or yellow? (“color” task) and was followed by a 2s response period with a blank screen. This sequence was repeated twice within a block with the same task cue but different movies. The subjects responded by pressing one of two buttons on a response box for each alternative in a task. The mapping of the buttons to the response was switched for the second run to rule out the effects of specific motor responses on the observed neural activations. We used a *physics task* > *color task* contrast to functionally identify the fronto-parietal physics regions in each subject individually.

#### Stability experiment

In addition to the physics ROI localizer, each participant also performed 4 runs of the main experiment. In this experiment, subjects viewed a sequence of images while maintaining fixation on a red dot at the center of the image and performed a 1-back task. Each run of this experiment contained 3 rest/fixation blocks and 12 20-second stimulus blocks (2 blocks for each of the 6 experimental conditions). Each run began with a fixation-only block followed by a random ordering of the blocks corresponding to the 6 experimental conditions. This was followed by another fixation-only block and the 6 experimental condition blocks shown in the reverse order. Each run ended with another fixation-only block. Each image block contained 10 trials including 2 1-back trials. In each trial, an image was shown for 1.8s followed by 0.2s of fixation-only interval. Subjects were instructed to maintain fixation (confirmed using eye-tracker for 6 out of the 13 subjects) and respond by pressing a button on the response box whenever the same image repeated one after the other in the sequence.

#### Convolutional Neural Network (CNN) analysis

Activations from the fc1000 layer (fully connected layer just preceding the softmax layer) of a Resnet-50 model trained on ImageNet object recognition challenge were extracted for both stable and unstable conditions across Block Towers, Physical-Objects and Physical-People scenarios. A linear Support Vector Classifier (SVM) was trained to distinguish between stable and unstable conditions within each scenario using 4-fold cross-validation. To test the generalizability of the learned classifier to other scenarios, the SVM classifier was tested on stability detection in the remaining two scenarios. We replicated the results using fc8 layer activations in an ImageNet pre-trained VGG-16 network.

### Data acquisition

All imaging was performed on a Siemens 3T MAGNETOM Tim Trio scanner with a 32-channel head coil at the Athinoula A. Martinos Imaging Center at MIT. For each subject, a high-resolution T1-weighted anatomical image (MPRAGE: TR = 2.53 s; TE = 1.64, 3.44, 5.24, 7.04 ms; α = 7°; FOV = 220 mm; Matrix = 220 x 220; Slice thickness = 1 mm; 176 slices; Acceleration factor = 3; 32 reference lines) was collected in addition to whole-brain functional data using a T2*-weighted echo planar imaging pulse sequence (TR = 2 s; TE = 30 ms; α = 90°; FOV = 216 mm; Matrix = 108 x 108; Slice thickness = 2 mm; Voxel size = 2 x 2 mm in-plane; Slice gap = 0 mm; 69 slices).

### Eye movement recordings

Eye movement data was recorded from 6 of the 13 subjects during both the physics ROI localizer task and the stability experiment using the EyeLink 1000 Eye-Tracker (SR Research) inside the scanner. We could not collect eye movement data from other subjects due to technical difficulties. Eye tracking data was preprocessed and analyzed to confirm that eye movements could not explain differences in BOLD activity for various experimental conditions. For each trial in both the localizer and stability tasks, we computed the average abscissa and ordinate of the eye position, the number of saccades, average duration and amplitude of saccades for the duration of the trial. We then performed t-tests to compare the average values of the aforementioned eye movement variables for stable and unstable conditions in each scenario across subjects.

### fMRI data preprocessing

Preprocessing was done using FreeSurfer (freesurfer.net). All other analyses were performed in MATLAB 2015B (The Mathworks). fMRI data preprocessing included motion correction, slice time correction, linear fit to detrend the time series, and spatial smoothing with a Gaussian kernel (FWHM = 5 mm). Before smoothing the functional data, all functional runs were co-registered to the subject”s T1-weighted anatomical image. All analyses were performed in each subject”s native volume and in some cases the results were plotted on the subject”s native inflated cortical surface only for better visualization (using FreeSurfer”s mri_vol2surf function). The general linear model included the experimental conditions and 6 nuisance regressors based on the motion estimates (x, y, and z translation; roll, pitch and yaw of rotation).

### Group-level physics parcel

We derived group-level physics parcels from the localizer data^16^ in 27 subjects using the Group-constrained Subject-Specific method described previously^45^. Briefly, individual subjects” binary activation maps (p < 0.005 uncorrected) were overlaid on top of each other in MNI space. This overlap map was spatially smoothed with a gaussian filter (FWHM = 8 mm) and then thresholded so that the map contained only those voxels with at least 10% overlap across subjects. Then, the overlap map was divided into group-level parcels using a water-shed image segmentation algorithm (*watershed* function in MATLAB). Finally, we selected a subset of parcels in which at least 16 out of the 27 (∼60%) subjects show some activated voxels. This resulted in 7 group-level parcels spanning frontal, parietal and occipital lobes. We rejected 2 of the 7 parcels in the occipital cortex since they were shown to respond to both physical and social stimuli^15^. The remaining parcels correspond coarsely to previously described physics regions^15,16^, however, we believe that the new parcels are probably more stable because they are derived from a larger subject pool. These five parcels were then combined to get one group-level parcel each in left and right hemispheres. We will make the parcels publicly available.

### Functional ROI definition

We defined functional regions of interest (fROI) in each individual subject as the intersection of subject specific localizer contrast map and group-level (or anatomical) parcels. Specifically, we used the physics localizer to identify brain regions in each individual subject that responded more to the physics task compared to the color task (uncorrected p-value < 0.001 for the physics > color contrast). This contrast map was then intersected with the group-level physics parcels created from the physics localizer data collected in a previous study^16^. Thus, the individual subject fROI contained only those voxels that showed significantly stronger activations for the physics task compared to the color task and fell within the group-level physics parcel. This allowed the fROI locations and sizes to vary across subjects but restricted them to a common general region across subjects. In addition to the physics fROI, we also defined fROIs for the primary visual cortex (V1), Lateral Occipital Complex (LOC) and Ventral Temporal Cortex (VTC) in each subject. As before, we used data from the physics localizer to identify brain regions that responded to visual stimuli compared to fixation (uncorrected p-value < 0.001 for physics + color > fixation contrast). We then intersected this contrast map with masks derived from anatomical parcellation and considered only those significant voxels from the contrast map lying within the anatomical mask for further analyses.

### Multi-voxel pattern correlation analysis

#### Within category separability

To assess if the candidate physics fROI holds information about physical stability, we used the multi-voxel pattern correlation analysis^20^. In an fROI, we computed *within* condition pattern correlations (correlation between voxel activation patterns for even and odd runs, computed separately for unstable and stable conditions within a scenario) and *between* conditions pattern correlations (correlation between voxel activation patterns for even runs of unstable condition and odd runs of stable condition within a scenario, and vice-versa). We computed the *within* and *between* condition pattern correlations for each scenario in each hemisphere in each subject and compared the magnitudes of correlations using appropriate statistical tests after transforming the correlations using Fisher transform (*atanh* function in MATLAB). A significantly higher *within* condition pattern correlation compared to *between* condition pattern correlation indicates that unstable and stable conditions evoke distinctive voxel activation patterns in a given fROI. Note that by always comparing data across even and odd runs we avoid obscuring pattern information from temporal pattern drifts^46^.

#### Across category similarity

To explore the generalizability of neural representation of physical stability in a given fROI, we computed multi-voxel pattern correlations computed across scenarios. Specifically, in a given fROI, we extracted activation patterns for unstable and stable conditions from the two scenarios under consideration (say, “Physical-Objects” and “Physical-People” scenarios). We then computed pattern correlations between unstable (or stable) conditions across scenarios (*within* condition), and pattern correlations between unstable and stable conditions across scenarios (*between* condition). In this way, we computed the four pairwise pattern correlations for each pair of categories within a given fROI in each hemisphere of each individual subject and transformed them using Fisher transformation. We compared the magnitudes of *within* and *between* condition correlations for each pair of categories across subjects using a paired t-test.

