## Supplementary Material for "Invariant representation of physical stability in the human brain"

| | | X location *(px)* | Y location *(px)* | Number of eye movements | Duration ofeye movements *(ms)* | Amplitude of eye movements *(minutes of arc)* | Interestingness rating *(scale 1-5)* |
|---|---|---|---|---|---|---|---|
| | | Screen resolution: 1024 x 768 (512, 384) is the fixation point | | | | | |
| Physical-Objects | unstable | 518.1 | 388.5 | 36.6 | 40.6 | 1.13 | **2.9** |
| | stable | 505.6 | 380.6 | 29.1 | 47.9 | 1.14 | **1.29** |
| | | *p = 0.12* | *p = 0.19* | *p = 0.1* | *p = 1* | *p = 0.93* | ***p = 0.0007*** |
| Physical-People | unstable | 517.8 | 377.2 | 37.7 | 30.4 | **1.31** | **3.69** |
| | stable | 516.9 | 370.8 | 31.9 | 39.9 | **1.08** | **1.97** |
| | | *p = 0.81* | *p = 0.6* | *p = 0.13* | *p = 0.62* | ***p = 0.028*** | ***p < 0.0001*** |
| Animals-People | unstable | 507.6 | 388.9 | 40.8 | 57.3 | 1.38 | **3.84** |
| | stable | 512.1 | 379.7 | 43 | 41.1 | 1.26 | **2.7** |
| | | *p = 0.81* | *p = 0.19* | *p = 0.45* | *p = 0.31* | *p = 0.26* | ***p = 0.0006*** |

Supplementary Table 1: Average values of eye tracking variables and interestingness ratings for stable and unstable conditions in all three scenarios. Each cell shows the average value of the variable for stable and unstable conditions along with the p-value for a paired t-test comparing the two sets of values across subjects. The first five columns correspond to eye movement variables collected on 6 subjects during the fMRI experiment and the last column is for the interestingness rating collected on the same set of images but in 11 subjects outside the scanner (see Methods for details). Since the subjects were instructed to maintain fixation at the center of the image, we did not observe any saccadic events (amplitude > 1 degree) and hence we are calling the small ballistic events as simply *eye movements*. Significant effects ($p < 0.05$) are highlighted in bold.

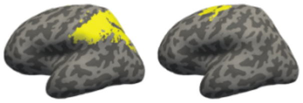

| Condition | Comparison | Parietal Physics ROI (# voxels: mean ± sem 273 ± 57.9) | Frontal Physics ROI (# voxels: mean ± sem 64.2 ± 13.6) |
|---|---|---|---|
| Physical-Objects | within | **0.62** | 0.40 |
| | between | **0.49** | 0.35 |
| | | ***p = 0.011*** | *p = 0.23* |
| Physical-People | within | **0.57** | 0.31 |
| | between | **0.49** | 0.23 |
| | | ***p = 0.002*** | *p = 0.089* |
| Animals-People | within | 0.42 | 0.34 |
| | between | 0.50 | 0.44 |
| | | *p = 0.013* | *p = 0.064* |
| Physical-People ↔ Physical-Objects | within | 1.13 | 1.25 |
| | between | 1.02 | 1.16 |
| | | *p = 0.077* | *p = 0.19* |
| Animals-People ↔ Physical-People | within | 1.20 | 1.28 |
| | between | 1.12 | 1.17 |
| | | *p = 0.18* | *p = 0.16* |
| Physical-Objects ↔ Animals-People | within | 1.06 | 1.24 |
| | between | 0.99 | 1.27 |
| | | *p = 0.37* | *p = 0.77* |

Supplementary Table 2: MVPA in the candidate physics regions done separately within parietal and frontal parcels. Each cell shows the average Fisher transformed within and between condition pattern correlations along with the p-value for a paired t-test comparing the two sets of values. Each column includes the results from one fROI. The top three rows contain results for within scenario pattern correlation analysis and the bottom three rows show results for the pattern correlation analysis across scenarios. Significant effects with within condition correlations greater than between condition correlations are highlighted in bold.

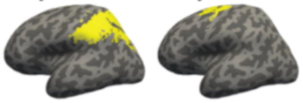

| Average beta estimates | | Parietal Physics ROI<br># voxels: mean ± sem<br>273 ± 57.9 | Frontal Physics ROI<br># voxels: mean ± sem<br>64.2 ± 13.6 |
|---|---|---|---|
| Physical-Objects | unstable | **0.16** | 1.75 |
| | stable | **-0.03** | 1.8 |
| | | ***p = 0.02*** | *p = 0.43* |
| Physical-People | unstable | **0.21** | 1.93 |
| | stable | **0.03** | 1.95 |
| | | ***p = 0.04*** | *p = 0.7* |
| Animals-People | unstable | 0.075 | 2.08 |
| | stable | 0.095 | 2.2 |
| | | *p = 0.67* | *p = 0.09* |

Supplementary Table 3: Average beta values for unstable and stable conditions in each of the scenarios computed separately for parietal and frontal physics parcels. Each cell shows average GLM estimated beta values for unstable and stable conditions along with the p-value for a paired t-test comparing the two sets of values across subjects. Scenarios showing significantly higher response to unstable scenes compared to stable scenes are highlighted in bold in each column.
